# Adenine base editing correction of *LMNA* c.745C>T (p.R249W) in congenital muscular dystrophy myoblasts improves cellular phenotype while revealing deleterious p.L248P bystander effects

**DOI:** 10.64898/2026.08.03.742539

**Authors:** M. Santafé, I. Hernández, D. Mazzeo, D. Gómez-Domínguez, D. Megías, K Mamchaoui, I. Pérez de Castro

## Abstract

**Background:** *LMNA*-related congenital muscular dystrophy (L-CMD) is a rare, life-threatening genetic disorder caused by point mutations in the *LMNA* gene, for which no effective treatment currently exists. It is characterized by early-onset muscle weakness, dropped-head syndrome, hypotonia, cardiac complications, and restrictive lung disease, frequently leading to premature death. The *LMNA* c.745C>T (p.R249W) mutation is the most prevalent amongst L-CMD patients. Given its monogenic nature, L-CMD represents a compelling candidate for gene therapy approaches.

**Results:** In this study, we investigated the therapeutic potential of adenine base editing (ABE) to correct the pathogenic *LMNA* c.745C>T (p.R249W) mutation in human myoblasts. We evaluated multiple ABE variants and single-guide RNAs (sgRNAs), identifying optimal combinations that achieved efficient and specific correction of the mutant allele. However, we found that editing can also introduce an adjacent bystander mutation, c.743T>C (p.L248P). To determine the functional consequences of base editing, we established clonal cell lines reverted to wild type or harboring the p.L248P variant. Whereas wild-type edited cells showed a clear correction for all the studied parameters that were abnormal in R249W myoblasts, we found that L248P cells show nuclear abnormalities resembling those of R249W mutant cells, and their cellular function is partially compromised. These results demonstrate that ABE can effectively target the *LMNA* c.745C>T mutation but also reveal the significant impact of bystander edits on cellular physiology.

**Conclusions:** Our findings provide proof-of-concept for the application of base editing as a therapeutic strategy for L-CMD, while underscoring the necessity of precise editing technologies to ensure both efficacy and safety in future clinical translation.

## BACKGROUND

The nuclear lamina is a filamentous structure underlying the inner nuclear membrane that provides mechanical support and organizes nuclear architecture (de Leeuw et al., 2018). In addition to preserving structural integrity, it regulates key cellular processes including DNA replication, transcription, chromatin organization, cell cycle progression, and differentiation (Busch et al., 2009; Dechat et al., 2008; Gruenbaum & Foisner, 2015; Reddy et al., 2008; Swift & Discher, 2014). In mammalian cells, the lamina mainly consists of type A lamins (lamins A and C, encoded by *LMNA*) and type B lamins (lamins B1 and B2, encoded by *LMNB1* and *LMNB2*) (Worman, 2012). Alterations in the nuclear lamina cause laminopathies, a broad spectrum of rare disorders, mostly caused by mutations in *LMNA*, that affect striated and cardiac muscle, adipose tissue, peripheral nerves, or multiple organs (Worman, 2012). To date, nearly five hundred pathogenic variants have been identified, resulting in at least 15 clinically distinct syndromes with variable severity (http://www.umd.be/LMNA/).

Among laminopathies, *LMNA*-related congenital muscular dystrophy (ORPHA:157973) represents one of the most devastating phenotypes. This autosomal dominant muscular dystrophy arises predominantly from *de novo* dominant negative mutations in *LMNA* and manifests within the first two years of life with hypotonia, dropped-head syndrome, progressive muscle weakness, early respiratory failure, and cardiomyopathy, leading to premature death during childhood (Ben Yaou et al., 2021). The missense mutation c.745C>T (p.R249W) is the most frequent cause of L-CMD (Ben Yaou et al., 2021; Cesar, Coll, et al., 2023; Fan et al., 2021) and results, at the cellular level, in a dysfunctional lamin A/C protein characterized by abnormal nuclear morphology, impaired mechanosensing, and accumulation of DNA damage (Bertrand et al., 2014, 2020; Earle et al., 2020; Gómez-Domínguez et al., 2020; Steele-Stallard et al., 2018). Currently, there is no cure for L-CMD, and clinical management is primarily supportive, aiming to alleviate symptoms and preserve function. Standard care includes physiotherapy and exercise programs, the use of assistive devices, surgical interventions when indicated, and regular monitoring of cardiac and respiratory function (Cesar, Campuzano, et al., 2023; Macquart et al., 2016). Given its severe clinical course, extremely low prevalence (<1 per 1,000,000) (https://www.orpha.net/en/disease/detail/157973), and lack of effective therapies, L-CMD remains a clear unmet medical need.

The precision of genome editing technologies has generated strong interest in their therapeutic application for rare diseases, including laminopathies. Classical CRISPR/Cas9 has shown promise in preclinical models for laminopathies and related disorders, including Hutchinson–Gilford progeria syndrome (Beyret et al., 2019; Santiago-Fernández et al., 2019), Duchenne muscular dystrophy (Agrawal et al., 2023) and L-CMD itself (Gómez-Domínguez et al., 2026). However, conventional Cas9 activity induces double-strand breaks and relies on error-prone repair pathways, making it poorly suited for correcting pathogenic single-nucleotide substitutions such as *LMNA* c.745C>T. By contrast, base editors represent a new generation of genome editing tools that enable direct and programmable conversion of single nucleotides without requiring double-strand breaks or donor templates (N. M. Gaudelli et al., 2017; Komor et al., 2016b; Nishida et al., 2016). ABEs (N. Gaudelli et al., 2017), in particular, enable A-T-to-G-C conversions and therefore provide a rational strategy to revert the *LMNA* c.745C>T mutation to wild type.

ABEs act by targeting a specific DNA sequence through a sgRNA and catalyzing a precise chemical conversion of one base to another within a defined editing window (Newby & Liu, 2021). This window is a limited stretch of nucleotides adjacent to the protospacer adjacent motif (PAM) where the editing enzyme can deaminate target bases. However, because the editing activity occurs within this window, nearby nucleotides can also be unintentionally modified, leading to so-called bystander mutations. Therefore, testing different ABE variants with distinct editing windows and constraints is critical for anticipating and minimizing unintended bystander base conversions and improving the precision of gene correction (Huang et al., 2021).

In this study, we explored adenine base editing as a therapeutic strategy for L-CMD. Specifically, we evaluated different ABE variants and sgRNAs designed to target the *LMNA* c.745C>T mutation in human myoblasts. We assess their efficiency, bystander activity, and phenotypic impact, focusing on nuclear morphology and other cellular features associated with lamin A/C dysfunction. This work establishes proof of principle for precise base editing in L-CMD patient-derived cells and provides a foundation for future translational approaches to treat *LMNA*-associated congenital dystrophies.

## RESULTS

### Analysis of ABE-Mediated correction of the *LMNA* c.745C>T (p.R249W) mutation in human myoblasts

Adenine base editors were used to revert to wild type the *LMNA* c.745T mutation causally associated with L-CMD. To minimize the chances of bystander generation (c.743T>C), six ABEs and five different sgRNAs were designed and tested to specifically target the *LMNA* c.745C>T (p.R249W) mutation in patient-derived human myoblasts (**Fig. 1** and **Table 1**). The sgRNAs were selected as optimally as possible to position the target adenine within the editing window while minimizing the risk of bystander mutations (**Fig. 1**). To shift the editing window and better center it on the target adenine while minimizing the susceptibility of adjacent adenines to unintended editing, extended versions of some selected sgRNAs were designed. For the AbeMAX-NRTH editor, sg04 was lengthened to generate sg04/21 (21 nucleotides) and sg04/22 (22 nucleotides) (**Fig. 1**). Similarly, for the Nme2 compact editor, sg06 and sg07 were extended to produce sg06/25 (25 nucleotides) and sg07/25 (25 nucleotides), respectively (**Fig. 1**).

**Figure 1.**
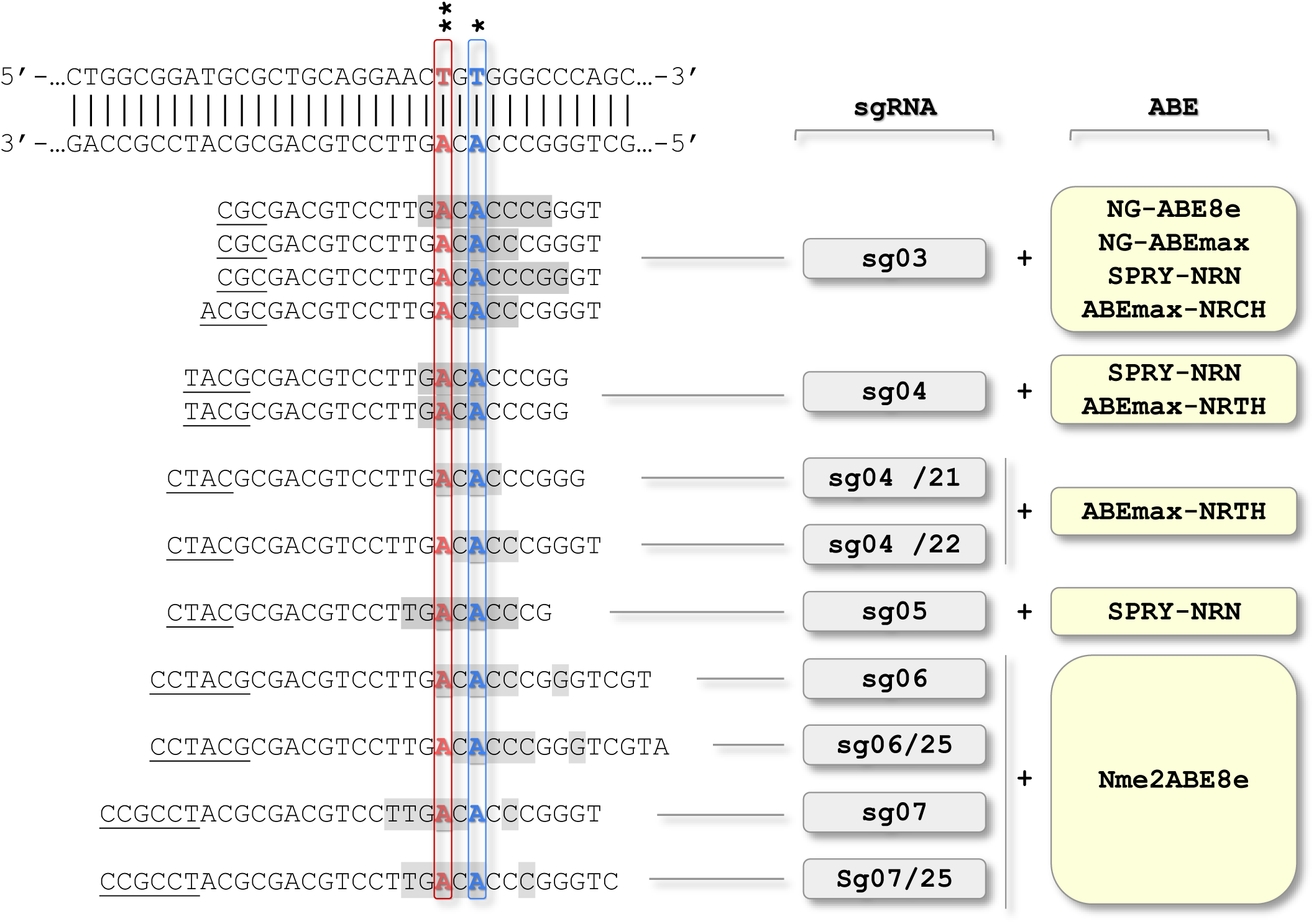
Adenine base editors and sgRNAs used for the correction of the *LMNA* c.745T mutation. Schematic representation of sgRNAs targeting the *LMNA* exon 4 mutation c.745C>T (p.R249W) together with the adenine base editors employed. The target adenine at position 745 is shown in blue (*), the potential bystander adenine at position 743 in red (**), the corresponding PAM sequences are underlines, and the predicted activity windows for each ABE–sgRNA pair are indicated in grey.

**Table 1.**
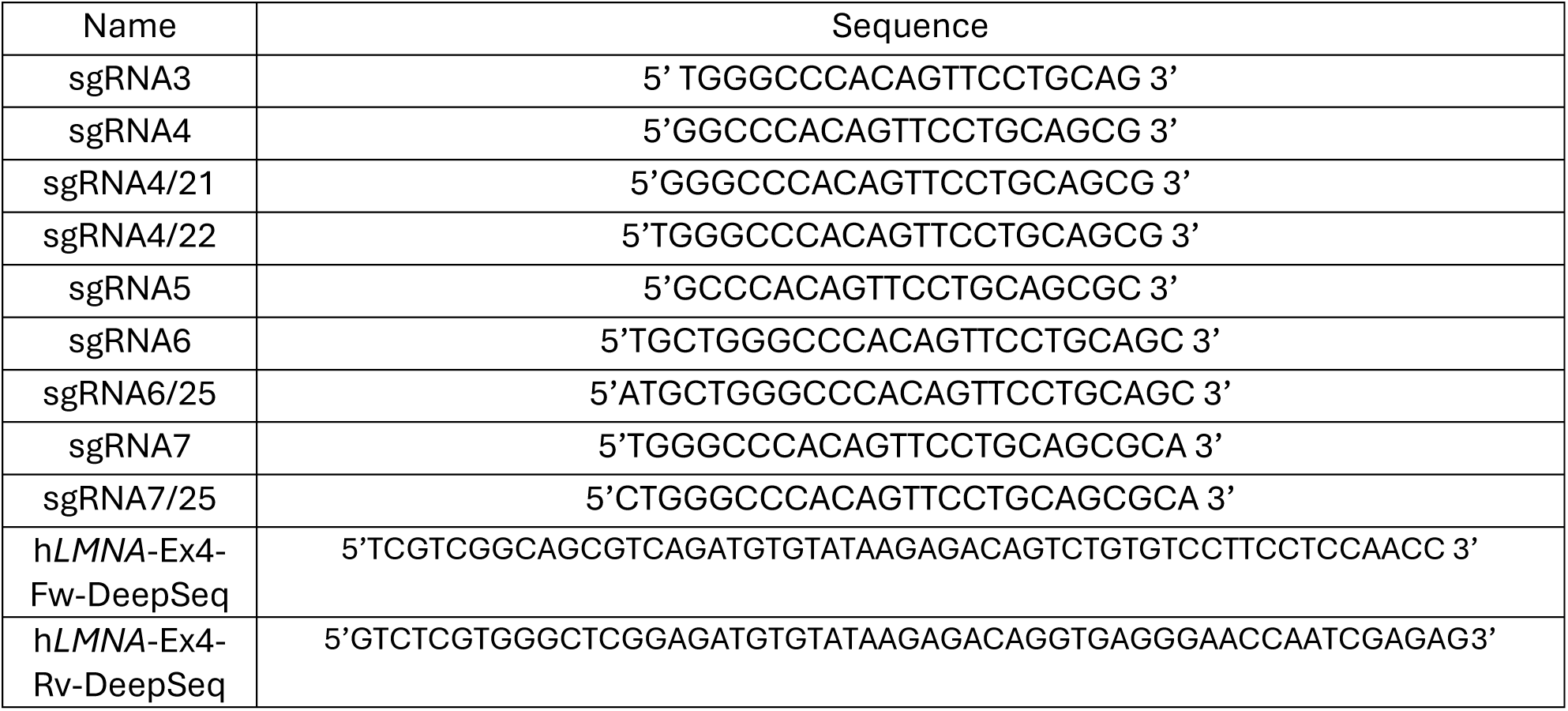
Sequence of sgRNAs and primers used in this study.

The panel of thirteen ABE-sgRNA combinations were co-delivered to the *LMNA* c.745T, patient cells and processed according to the work plan depicted in **Figure 2A**. Cotransfection of a GFP reporter plasmid allowed enrichment of transfected cells by fluorescence-activated cell sorting. As a negative control, cells were transfected with ABE constructs in the absence of sgRNAs. Efficient nucleofection of ABE-sgRNA complexes was achieved, with GFP-positive cells ranging between 8% and 20% after 24 hours (**Fig. S1**).

**Figure 2.**
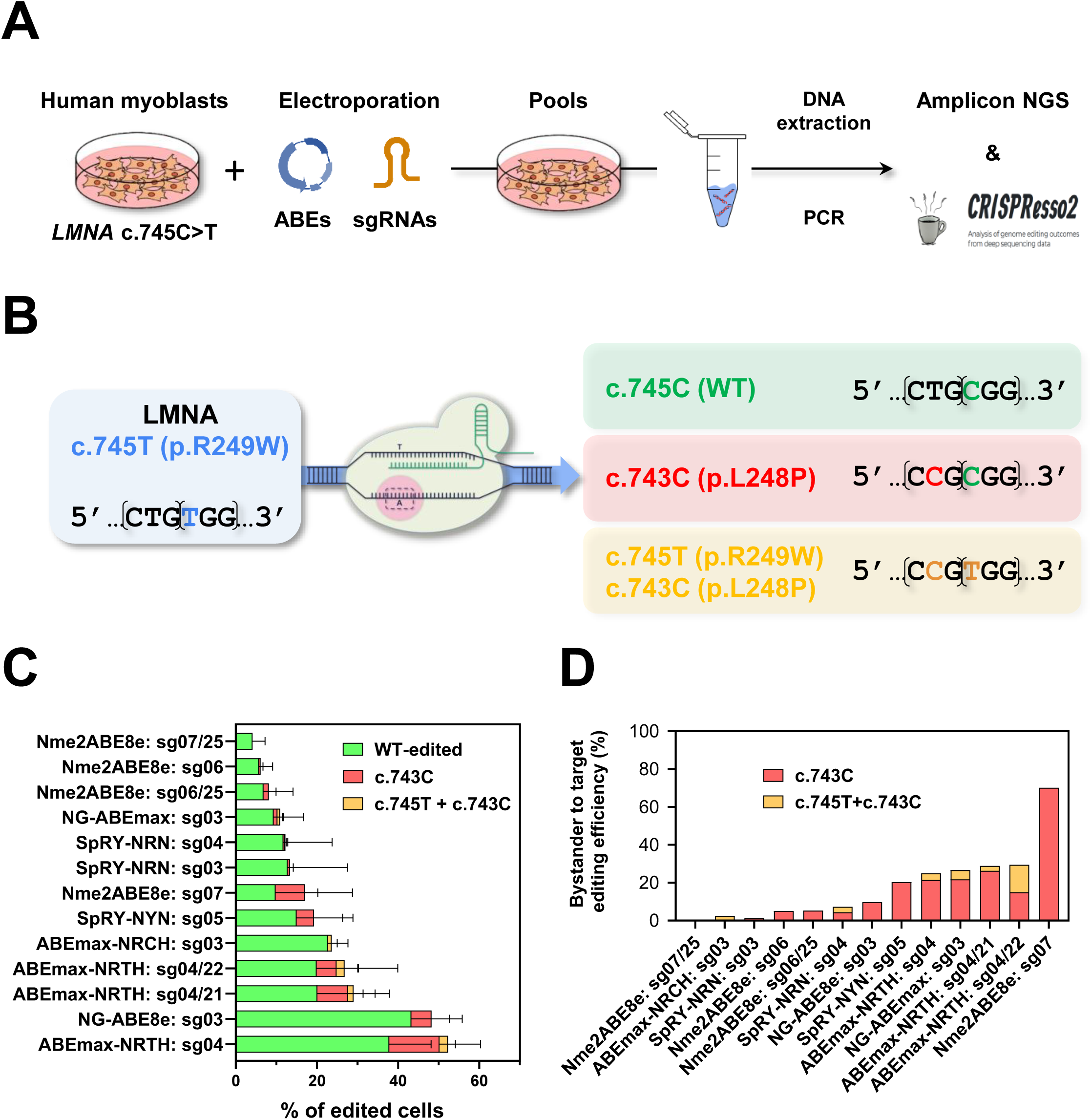
Adenine base editing corrects *LMNA* c.745C>T in human myoblasts with variable efficiency and minimal L248P bystanders. **(A)** Schematic experimental design for the assessment of ABE/sgRNA activity in human myoblasts. Various biological replicates have been performed (n = 2–4 independent pools per condition). **(B)** Graphical representation of ABE mechanism of action over mutation *LMNA* c.745C>T and the expected alleles to be found upon gene editing. **(C)** Percentage of reads obtained for the ABE-sgRNA combinations for each edited allele was used to estimate the percentage of all the possible edited cell types: WT-edited, c.743C (carrying a bystander mutation), c.743C+c.745T (carrying bystander and parental mutations). **(D)** Analysis of the ratio of unintended bystander nucleotide edits relative to the intended on-target edit for different adenine base editor (ABE) and sgRNA combinations. A lower ratio indicates higher precision with fewer bystander edits.

Genomic DNA was isolated from nucleofected cell pools, and editing efficiencies were quantified by next-generation sequencing (NGS) of PCR-amplified fragments spanning the target region. Given the neighboring thymine at position c.743, base editing was predicted to potentially introduce a bystander substitution c.743T>C (p.L248P) together with or independent of target correction. Consistent with this, NGS analysis identified four major allelic classes in edited populations: (i) unedited alleles retaining c.745C>T (known as parental or p.R249W from now on), (ii) wild-type alleles (endogenous WT or corrected; known as WT-edited from now on), (iii) alleles carrying only the bystander substitution (known as bystander or p.L248P from now on), and (iv) alleles harboring both p.R249W mutation and bystander editing (**Fig. 2B**). The distribution of allele outcomes after sequencing varied depending on the specific combination of ABE and sgRNA used (**Fig. S2**) and was used to estimate the percentage of cells carrying each of the edited configurations (**Fig. 2C**). In control samples, allele distribution averaged 50% wild-type and 50% mutant c.745C>T sequences, reflecting the heterozygous background of the parental cell line (**Fig. S2**). The highest edition level was achieved with ABEmax-NRTH and sg04, reaching 52.29% of overall editing and 37.83% precise reversion to wild-type without bystander generation (**Fig. 2C**; **Fig. S2**). ABE8e with sg03 also displayed robust activity, with 48.2% overall editing and the highest 43.3% exact correction (**Fig. 2C**; **Fig. S2**). Additional conditions achieved moderate efficiencies (10-30%) with also a variable on-target correction efficiency (**Fig. 2C**; **Fig. S2**). Three combinations yielded <10% correction, all with compact editor Nme2ABE8e, namely Nme2ABE8e:sg06/25, Nme2ABE8e:sg06, and Nme2ABE8e:sg07/25 with editing efficiencies of 8.13%, 6.11% and 4.15% respectively.

The percentage of bystander editing introduced by each on-target mutation was calculated to assess the extent of unintended base conversions within the editing window (**Fig2.D**). Bystander analysis revealed the highest levels of the c.743T>C (p.L248P) allele in condition Nme2ABE8e:sg07 (70.16%) (**Fig. 2D**). Importantly, seven combinations resulted in <10% of bystander alleles per corrected event which from highest to lowest were ABE8e:sg03 (9.74%), SpRY-NRN:sg04 (7.26%), Nme2ABE8e:sg06/25 (5.28%), Nme2ABE8e:sg06 (5.1%), ABEmax-NRCH:sg03 (2,5%), SpRY-NRN:sg03 (1.23%) and Nme2ABE8e:sg07/25 (0%) (**Fig.2D**), suggesting that careful selection of guide-editor pairs can minimize bystander editing at position c.743.

ABEmax-NRTH editor with its longer versions of sg04/21 and sg04/22 showed a slight reduction in the bystander edition but accompanied with a parallel reduction of the overall editing activity, which implies that these extra nucleotides do not improve neither activity nor specificity of the editor (**Fig. 2C-D**). A similar pattern was seen with editor Nme2ABE8e and sg06/25 (**Fig 2C-D**). However, with sg07/25 compared to its normal length version, a drastic reduction was seen in the introduced bystander allele (**Fig 2C-D**).

In summary, base editing using selected ABE–sgRNA combinations effectively corrected the *LMNA* c.745C>T mutation in human myoblasts with efficiencies ranging from 10% to 50%. While bystander editing at position c.743T>C occurred under some conditions, its frequency was generally low, underscoring the potential of optimized ABE approaches for therapeutic correction of *LMNA*-associated muscular dystrophies.

### Nuclear morphology evaluation in edited and non-edited p. R249W myoblast clones

The therapeutic potential and efficacy of ABEs for L-CMD were further assessed by phenotypically characterizing the wild-type corrected myoblast clones derived following genome editing. To this end, several cellular features, previously shown to be disrupted in *LMNA* c.745C>T (p.R249W) myoblasts, were systematically evaluated in corrected myoblasts (**Fig. 3A**). The phenotype of cells harboring one copy of the bystander *LMNA* c.743T>C (p.L248P) modification was also investigated within the same experimental framework (**Fig. 3A**).

**Figure 3.**
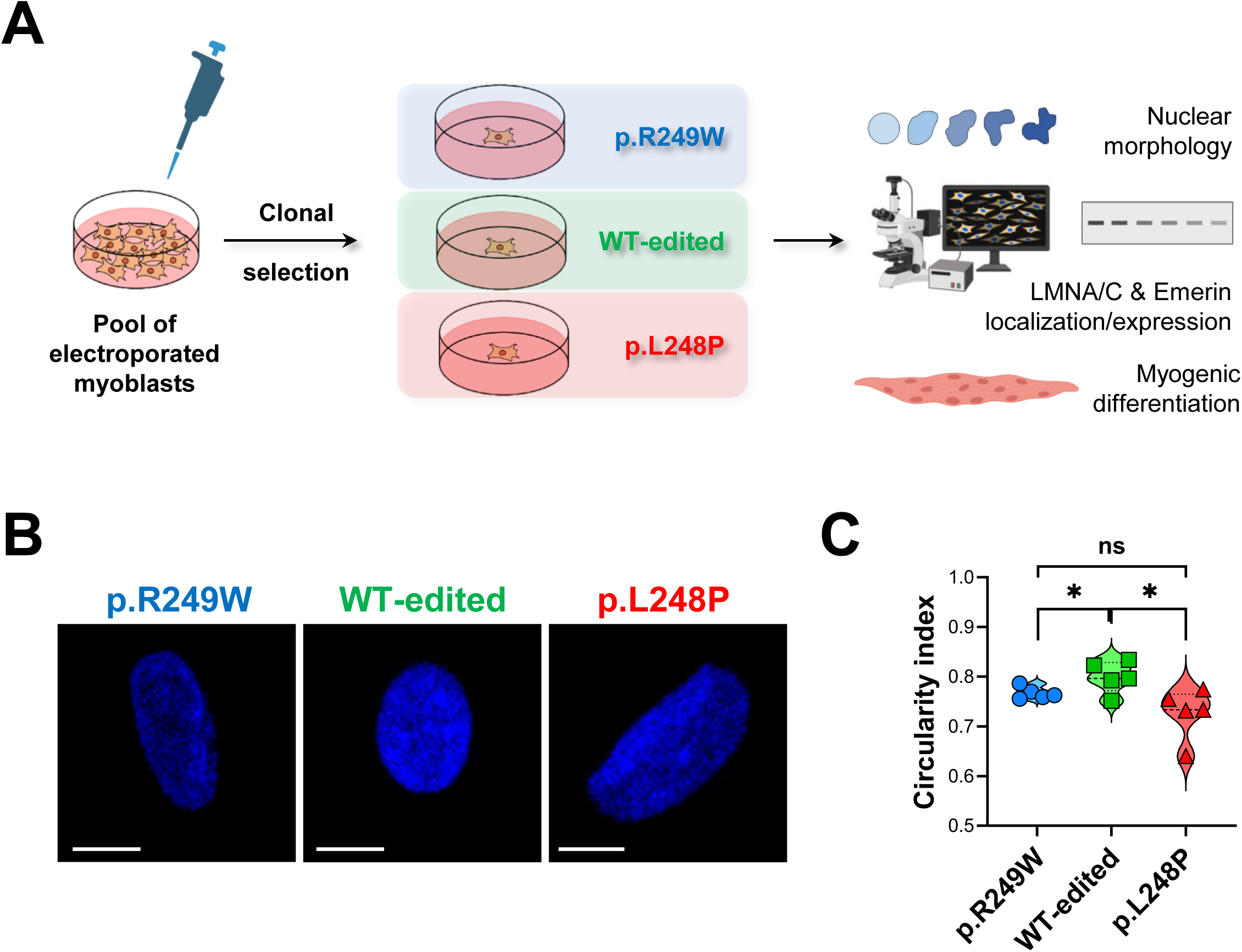
Nuclear morphology improves after *LMNA* c.745T base editing, but L248P bystander clones retain mutant-like morphology. **(A)** Schematic experimental design for generation and characterization of WT-edited (c.745T>C), p.R249W (c.745T, parental) and p.L248P (c.743C, bystander) human clones. **(B)** Representative microscopy images from nuclei for each clone type stained with Hoechst. **(C)** Circularity index was calculated in nuclei (n>50 nuclei per clone) from clones of the three genotypes growing in 2D cultures (n=5-6 clones per genotype). Scale bar = 10 μm. One tailed, unpaired t-test analysis was performed for the statistical analysis, where ns p>0.05; * p<0.05; ** p<0.01; *** p<0.001. Data is presented as the average values of each clone obtained in a single experiment. Graph represents one of the two biological replicates carried out.

Editing the *LMNA* c.745C>T mutation to the wild-type sequence in human myoblasts using ABEs resulted in a significant improvement in nuclear circularity compared to the parental mutant cells (**Fig. 3B-C**). This finding is consistent with previous studies reporting compromised nuclear morphology and reduced circularity associated with the p.R249W mutation in myoblasts (Gómez-Domínguez et al., 2026; Steele-Stallard et al., 2018).However, clones carrying the bystander c.743T>C (p.L248P) mutation exhibited no differences in nuclear circularity relative to p.R249W clones (**Fig. 3B-C**). This observation suggests that the bystander mutation negatively impacts nuclear morphology, potentially leading to a phenotype similar to the pathogenic p.R249W mutant. Overall, these results support the therapeutic benefit of precise correction of the p.R249W *LMNA* mutation and suggest a pathogenic potential of the p.L248P variant.

### Lamin A/C and Emerin expression and nuclear envelope localization is improved in p.R249W myoblast clones edited to wild type

Mutations in *LMNA* have been associated with a reduced expression of LMNA/C proteins (Gómez-Domínguez et al., 2020; Steele-Stallard et al., 2018). To evaluate the impact of ABE-mediated editing on this phenotype, we assessed protein expression by Western blot. WT-edited clones exhibited increased expression levels of lamin A/C both individually and combined (**Fig. 4A**), compared to parental p.R249W mutant and bystander clones. Additionally, bystander clones did not show significant differences compared to the p.R249W mutants (**Fig. 4A**), suggesting that the mutant phenotype characteristics are maintained in these bystander-edited cells.

**Figure 4.**
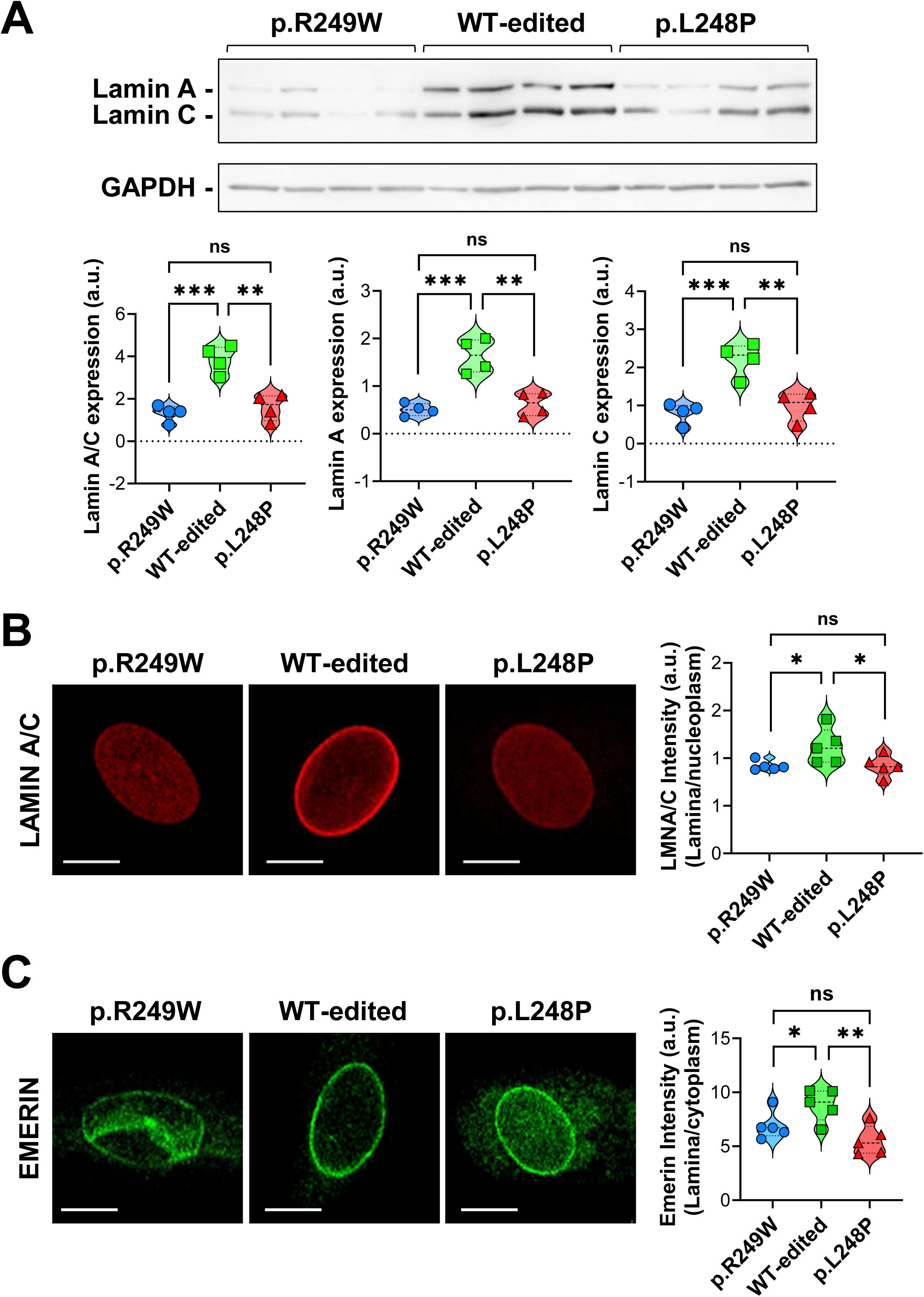
ABE-corrected *LMNA* p.R249W myoblasts show increased lamin A and C expression and improved lamin/emerin localization, whereas p.L248P clones retain mutant phenotype. **(A)** Representative immunoblot for the detection of lamin A/C in WT-edited, p.R249W and p.L248P clones (n=4 clones per genotype). This represents one of the three technical replicates carried out. Bottom graphs show the quantification of lamin A/C expression in WT-edited, p.R249W and p.L248P clones. **(B)** WT-edited clones are characterized by an increased ratio of lamina/nucleoplasm lamin A/C expression when compared with parental (p.R249W) and bystander (p.L248P) clones. Microscopy images show representative single lamin A/C nuclei for each clone type. Graph shows average fluorescence intensity calculated for lamin A/C signal at lamina and normalized with the nucleoplasm signal (n ≥ 100 nuclei per clone) (n=5-6 clones per genotype). **(C)** Cytoplasmic signal of Emerin is reverted in WT-edited clones when compared with p.R249W and p.L248P clones. Representative single Emerin microscopy images from nuclei are shown for each clone type. Graph shows the average fluorescence intensity calculated as the ratio between the nuclear and the cytoplasmic levels (n ≥ 100 nuclei per clone) (n=5-6 clones per genotype). Scale bar = 10 μm. In all cases one tailed, unpaired t-test analysis was performed for the statistical analysis of clone-type comparisons, where ns p>0.05; * p<0.05; ** p<0.01; *** p<0.001. Data is presented as the average values of each clone obtained in a single experiment. This represents one of the two biological replicates carried out.

Additionally, mutations in *LMNA* can impair the integrity of the nuclear lamina and disrupt the distribution of key inner nuclear membrane proteins. Hence, increased lamin A/C expression at the nucleoplasm has been reported in myoblasts carrying LMNA mutations (Bertrand et al., 2020). Furthermore, Emerin, a critical component of the nuclear lamin, mislocalize to the cytoplasm in L-CMD R249W myoblasts (Storey et al., 2024). Subsequently, confocal imaging was performed to analyze the subcellular localization of lamin A/C and lamin-associated protein Emerin. Significant restoration of nuclear membrane localization was observed in WT-edited clones relative to parental, p.R249W, myoblasts (**Fig. 4B** for lamin A/C; **Fig. 4C** for Emerin). Notably, bystander clones did not differ significantly from parental cells (**Fig. 4B-C**), suggesting that correction to wild-type sequence enhances the proper nuclear localization phenotype, whereas introduction of the bystander mutation reverts localization to a state comparable to the p.R249W background.

### Myogenic differentiation improvement in base-edited, corrected myoblast clones

To analyze the differentiation potential of edited myoblasts, clones from the three groups (p.R249W, WT-edited, and p.L248P) were induced to form myogenic fibers and followed during 120 hours by time-lapse videomicroscopy tracking their nuclei (**Fig. 5A**). As expected for their pathogenic properties, nuclei from parental, p.R249W clones do not aggregate in a significant manner (**Fig. 5A**). However, nuclei from WT-edited cells move closer and form aggregates that increased in area with time (**Fig. 5A**). Interestingly, bystander clones displayed an intermediate phenotype (**Fig. 5A**).

**Figure 5.**
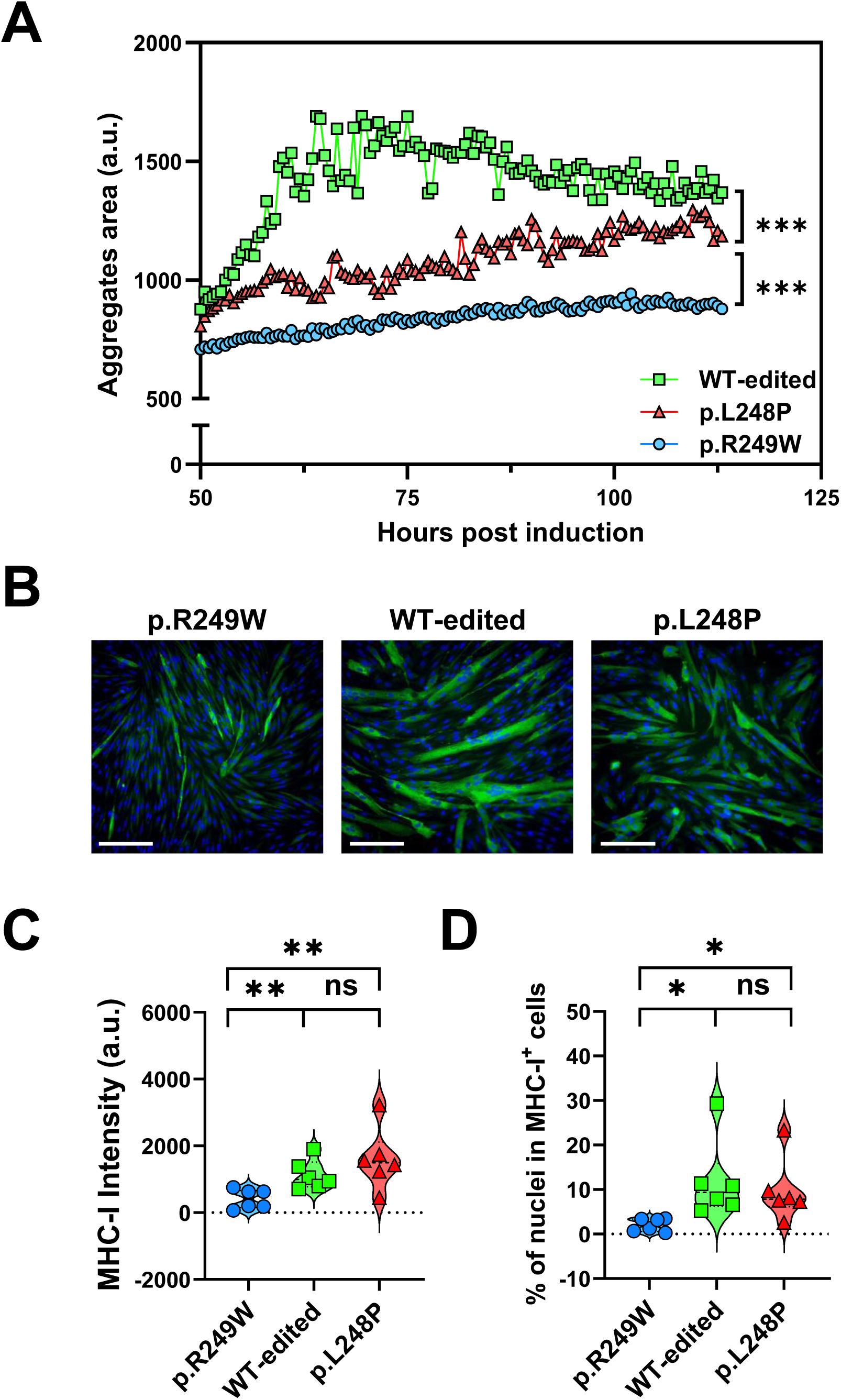
Myogenic differentiation is improved in ABE-corrected *LMNA* p.R249W myoblasts, whereas show intermediate phenotype in p.L248P, bystander clones. **(A)** Analysis of area of aggregates formed in WT-edited, p.R249W and p.L248P differentiating clones from 48 to 120 hours after differentiation induction. n=3 clones per genotype. **(B)** Representative single MHC-I confocal images for each *LMNA* group are shown at 72 hours post myogenic differentiation induction. Scale bar = 100 μm. **(C)** Analysis of MHC-I intensity normalized per nuclei of in WT-edited, p.R249W and p.L248P differentiating clones (n=6 clones per genotype). **(D)** Quantification of the percentage of nuclei in MHC-I positive cells in WT-edited, p.R249W and p.L248P differentiating clones (n=6 clones per genotype). One tailed, unpaired t-test analysis was performed for every statistical analysis, where ns p>0.05; * p<0.05; ** p<0.01; *** p<0.001. This represents one of the two biological replicates carried out.

To further explore myogenic formation capacity, myosin heavy chain 1 (MHC-I) expression was evaluated by immunofluorescence 72 hours after differentiation induction. This marker of myogenic differentiation was expressed at higher levels in WT-edited clones than in parental myoblasts, whereas no significant differences were observed when compared with bystander clones (**Fig. 5B-C**). Similar results were obtained when nuclei fusion was estimated in these differentiated samples. Hence, WT-edited clones showed a significant increase of nuclei included in MHC-I positive cells when compared with parental clones, whereas no significant differences were observed relative to bystander clones (**Fig. 5D**).

These data indicate that ABE-mediated editing to wild-type sequence enhances myogenic differentiation capacity of *LMNA* mutant myoblasts, as evidenced by increased aggregate formation, MHC-I expression and nuclear fusion. The bystander mutation confers partial phenotypic rescue with significant improvements in the analyzed differentiation metrics, suggesting an intermediate functional effect.

## DISCUSSION

Base editing is a precise and promising CRISPR-based therapeutic approach for genetic diseases caused by point mutations, offering correction without inducing double-stranded DNA breaks (N. Gaudelli et al., 2017; Komor et al., 2016a). In this study, we applied ABEs to correct the pathogenic *LMNA* c.745C>T (p.R249W) variant responsible for L-CMD. Through extensive screening of six ABE variants combined with nine sgRNAs (yielding thirteen editor–sgRNA combinations), we observed significant variability in editing efficiency and specificity. The highest overall editing was achieved using ABEmax-NRTH paired with sg04 (52.3%), closely matched by ABE8e:sg03, which showed the highest on-target correction efficiency. Contrary to some reports suggesting that extending sgRNA length improves editing specificity by better positioning the target nucleotide (Perrotta et al., 2024), our data using varying sgRNA lengths with the near-PAM-less ABEmax-NRTH editor indicated no differences in editing efficiency or specificity. The ratio of bystander to on-target editing was consistent across sgRNA lengths, suggesting that sgRNA extension beyond 20 nucleotides did not substantially modulate activity in this context.

We have also analyzed a compact base editor, Nme2ABE8e, advantageous for potential single-AAV delivery thanks to its limited size (Davis et al., 2022). As expected because of their previously reported activity (Edraki et al., 2019), this compact editor showed reduced editing efficiencies (<20%) relative to non-compact systems. Furthermore, mixed observations were made when longer sgRNAs were used, as the sg07/25 did reduce the appearance of the bystander edit highlighting possible editor and target-specific effects, whereas the sg06/25 increased editing activity without affecting bystander generation.

To understand the consequences of base editing at *LMNA* c.743T>C we have analyzed in detail myoblast clones reverted to wild type and those that, besides correcting this mutation, acquired a new one at an adjacent thymine (c.743T>C, p.L248P). To note, the bystander p.L248P variant has been listed as a variant associated with Emery-Dreifuss muscular dystrophy (EDMD) (Vytopil et al., 2003) but, to our knowledge, had not been functionally characterized. Phenotypically, WT-corrected clones displayed significant restoration of nuclear morphology, as evidenced by increased nuclear circularity, along with rescued expression and localization of lamin A/C and Emerin at the nuclear envelope and myogenic differentiation capacity. On the other hand, clones carrying one p.L248P bystander variant exhibited a mixed phenotype, with most parameters resembling p.R249W mutants, while others suggested partial functional recovery. This intermediate phenotype aligns with clinical insight that p.L248P is associated with EDMD, a milder condition than L-CMD. Because editing acts on the mutant allele while leaving the second allele wild type, the p.L248P bystander arises in heterozygosity, mirroring how it would occur in a treated patient. Therefore, our findings suggest that base editing correction of the *LMNA* c.745C>T variant may benefit L-CMD patients even if it induces the *LMNA* c.743T>C bystander mutation. Beyond this specific balance between on-target benefit and bystander risk, it is worth placing our approach within the broader landscape of gene-editing strategies for laminopathies.

Gene therapy has been increasingly explored for the treatment of laminopathies, a group of rare diseases caused by defects in nuclear lamina components. Most efforts have focused on Hutchinson-Gilford progeria syndrome (HGPS), where promising results have been obtained using both CRISPR 1.0 nuclease approaches (Beyret et al., 2019; Santiago-Fernández et al., 2019) and, more recently, base editors (Abutaleb et al., 2025; Gete et al., 2021; Koblan et al., 2021; Whisenant et al., 2022), a more precise CRISPR-based technology. Base editing has also been applied to correct *LMNA* mutations and partially revert pathogenic phenotypes in *LMNA*-related cardiac diseases (Caravia et al., 2025; Yang et al., 2024), and precise gene correction has been used in an L-CMD cell line as a readout of *LMNA*-regulated gene expression (Wang et al., 2022). Despite these advances, therapeutic options for L-CMD remain limited: to date, only spliceosome-mediated RNA trans-splicing has been tested as a preclinical strategy, with low-efficiency outcomes (Azibani et al., 2018), together with a CRISPR-Cas9 nuclease approach designed to selectively eliminate the mutant *LMNA* c.745C>T allele, which improved cardiac function and survival in a *Lmna^R24SW^* mouse model (Gómez-Domínguez et al., 2026). Such allele-elimination strategies, however, generate a hemizygous or null state rather than restoring the wild-type sequence. To our knowledge, the present study is the first to apply adenine base editing to precisely correct the most frequent L-CMD mutation (*LMNA* c.745C>T, p.R249W) in patient-derived myoblasts and to functionally characterize the resulting phenotypic rescue, spanning nuclear morphology, nuclear-envelope protein localization and myogenic differentiation. At the same time, our results illustrate a challenge specific to base editing at this locus, namely the generation of the p.L248P bystander, which allele-elimination approaches do not face.

Taken together, these results emphasize that combinatorial optimization of both the base editor and the guide RNA is critical to maximize on-target correction while minimizing bystander generation. Our study also has limitations. The phenotypic rescue was demonstrated in patient-derived myoblasts from a single L-CMD individual and in an *in vitro* setting, using plasmid electroporation as the delivery method. Validation in additional patient backgrounds and, ultimately, *in vivo* will be required to establish therapeutic efficacy. Addressing these aspects will be essential to determine whether adenine base editing, or alternative precise technologies such as prime editing (Anzalone et al., 2019), offers the most suitable strategy for *LMNA* c.745C>T-driven L-CMD.

## CONCLUSIONS

This study reports, for the first time, the successful use of adenine base editing (ABE) to revert the *LMNA* c.745C>T (p.R249W) pathogenic variant to the wild-type sequence in human myoblasts. Efficient reversion to wild-type was achieved in more than 30% of cells with four of the thirteen ABE/sgRNA combinations we have explored, with ABE8e– sg03 showing the highest efficiency. A bystander *LMNA c*.743T>C (p.L248P) variant emerged in a variable fraction of the corrected cells. This variant, previously associated with EDMD, has been characterized here for the first time. Importantly, ABE-mediated on-target correction improved nuclear morphology and Lamin A/C expression, and normalized Lamin A/C and Emerin subcellular localization. However, it produced a nuclear phenotype resembling that of the p.R249W mutation in p.L248P cells. Edited myoblasts also regained proper differentiation capacity, while bystander *LMNA* ^+/L248P^ cells exhibited a slight functional rescue.

In conclusion, this work establishes adenine base editing as a viable therapeutic strategy for *LMNA*-related muscular dystrophies. It illustrates the delicate balance required between editing efficiency, specificity, and phenotypic restoration to maximize patient benefit. Continued refinement in editor design, delivery, and safety evaluation will be essential to fully realize the clinical potential of base editing therapies.

## MATERIALS AND METHODS

All antibodies, plasmids, cell-culture reagents and media, equipment and software used in this study are listed in the Key Resources Table (Table S1).

### Base editors and sgRNAs

To correct the c.745C>T mutation, up to nine guide sgRNAs were designed with ABE constructs specific towards the mutant human *LMNA* allele (**Table 1**) using the Breaking-Cas design tool (Oliveros et al., 2016). With the main objective to locate the target mutation inside the editing window, each of them has a different PAM sequence. The base editors used were non-compact editors NG-ABE8e (Richter et al., 2020), NG-ABEmax (Huang et al., 2019), ABEmax-SpRY (Huang et al., 2023), ABEmax-NRCH (Miller et al., 2020) and ABEmax-NRTH (Miller et al., 2020); and compact editor Nme2ABE8e (Davis et al., 2022).

### Cell lines

The editing efficiencies of the ABEs were tested in an immortalized human myoblast line derived from a 3-year-old L-CMD patient carrying the *LMNA* c.745C>T (p.R249W) mutation (line EMD2784) obtained from the Institut de Myologie (Paris, France) and have been described previously (Bertrand et al., 2014; Gómez-Domínguez et al., 2020). The cells were cultured at 37 °C and 5% CO2. The culture medium was composed of one volume of Medium 199, 4 volumes of DMEM supplemented with 20% fetal bovine serum (FBS), 1× penicillin/streptomycin, 275 μg/ml insulin, 25 μg/ml fetuin, 0.5 ng/ml basic fibroblast growth factor (FGFb), 5 ng/ml human epidermal growth factor (EGF) and 0.2 μg/ml dexamethasone.

### Cell transfection and enrichment

Human myoblasts were transfected at 75–80% confluence by nucleofection on a NEPA21 Type II electroporator (Nepagene). One million cells per cuvette in 100 μl of Opti-MEM were electroporated with a poring pulse (275 V, 1 ms, 50 ms interval, 2 pulses, 10% decay, + polarity) followed by a transfer pulse (20 V, 50 ms, 50 ms interval, 2 pulses, 40% decay, ± polarity). Each condition received 10 μg of the corresponding ABE plasmid and 1 μg of pmaxFP-green (GFP), used to estimate transfection efficiency by fluorescence microscopy and to enable enrichment, together with 3 μM of sgRNA.

Twenty-four hours after transfection, GFP-positive cells were enriched by fluorescence-activated cell sorting. Nucleofected cells were sorted in PBS with 10% FBS, selecting GFP-positive cells as a proxy for uptake of the sgRNA and editing machinery, and the recovered pool was expanded in culture medium for DNA extraction.

### DNA sequencing and bioinformatic analysis

Editing was quantified by deep sequencing. After DNA extraction, *LMNA* exon 4 was amplified by PCR with specific primers (h*LMNA*-Ex4-Fw-DeepSeq and h*LMNA*-Ex4-Rv-DeepSeq; **Table 1**) carrying Illumina adapters, and sample indexes were added in a second amplification. PCR products were gel-purified, quantified by fluorometry, pooled, and sequenced on an Illumina NextSeq. FASTQ files were analyzed with CRISPResso2 (Clement et al., 2019) (https://crispresso.pinellolab.org/) to determine read counts and editing percentages for each condition.

### Single-cell cloning

Clones were generated from the pool previously edited with NG-ABE8e and sg03 by limiting dilution to one cell per well in 96-well plates. Grown clones were first screened by Sanger sequencing to identify unedited (parental) clones retaining the disease allele (*LMNA*^+/R249W^). Edited clones were then characterized in detail by deep sequencing and assigned to the two remaining groups: WT-edited (corrected to wild type at c.745; *LMNA*^+/+^) and bystander (corrected for the c.745C>T variant but carrying the c.743T>C p.L248P edit; *LMNA*^+/L248P^). The twenty-four clones used in this work were: parental (pH6-3, pE5-2, pC2-3, pH12-2, pC1-2 and pA6-3; n = 6); WT-edited (wA11-3, wA6-2, wC5-5, wC11-4, wF11-3, wC10-2, wF9-2, wA6-3 and wD2-3; n = 9); and bystander (bD2-3, bC6-5, bA9-3, bD6-5, bF9-2, bC9-5, bD7-3, bA11-3 and bF11-3; n = 9).

### Nuclear morphology analysis

Human myoblast clones were seeded on a μ-Slide 18-well plate (5,000 cells/well) and, at 50% confluence, fixed with cold methanol (−20 °C, 5 min). Cells were blocked with 100 μl/well of blocking solution (1% bovine serum albumin [BSA], 2.5% goat serum, 2.5% donkey serum and 0.3% Triton X-100 in PBS) and stained with Hoechst (1:5,000). Nuclear images were acquired on a Stellaris 8 Falcon STED confocal microscope (Leica Microsystems) with a 20× objective and processed in Arivis. The nucleus was selected as the region of interest (ROI), and the circularity index was determined with the corresponding software tool. More than 50 ROIs were analyzed per clone and averaged.

### Western blot analysis

Proteins were extracted from frozen cell pellets in RIPA buffer containing protease inhibitors, PMSF, orthovanadate and NaF. Samples were sonicated (Ultrasonic Processor UP100H, Hielscher; 3 × 5 s), incubated on ice for 10 min and centrifuged (13,000 rpm, 15 min, 4 °C) to pellet debris. Protein concentration was determined with the Pierce BCA Protein Assay. Equal amounts of protein were resolved by SDS-PAGE on Criterion TGX Stain-Free precast gels and transferred with the Trans-Blot Turbo system. Membranes were blocked in 5% milk in T-PBS and incubated with primary antibodies against lamin A/C (E-1; 1:3,000) and GAPDH (14C10; 1:2,000), followed by HRP-conjugated anti-mouse (NA931; 1:5,000) and anti-rabbit (NA934; 1:5,000) secondary antibodies. Immunodetection was performed with Pierce ECL substrate on an Amersham ImageQuant 800.

### Nuclear lamin A/C and Emerin immunofluorescence

Human myoblast clones were seeded on a μ-Slide 18-well plate (5,000 cells/well) and, at 50% confluence, fixed with cold methanol (−20 °C, 5 min). Cells were blocked with 100 μl/well of blocking solution (1% BSA, 2.5% goat serum, 2.5% donkey serum and 0.3% Triton X-100 in PBS) and incubated overnight at 4 °C with primary antibodies against lamin A/C (E-1; 1:500) and emerin (D3B8G; 1:1,000) in PBS. After three 5-min washes in PBS with 0.3% Triton X-100, cells were incubated with secondary antibodies (goat anti-mouse IgG, Alexa Fluor 594; goat anti-rabbit IgG, Alexa Fluor 488) for 1 h at room temperature in the dark. After two washes at 37 °C, images were acquired on a Stellaris 8 Falcon STED confocal microscope (Leica Microsystems) with a 20× objective. At least 50 nuclei per clone were analyzed, and maximum-intensity projections were used to quantify lamin A/C signal. Nuclei were selected as ROIs in Arivis, and the average ratios between nuclear-membrane, nucleoplasm and cytoplasm signals were calculated.

### Videomicroscopy of human differentiated myoblasts and bioinformatic quantification

Human myoblast clones were seeded on an 18-well plate (25,000 cells/well) in growth medium. At 80–90% confluence, the medium was replaced with 100 μl of differentiation medium (2% horse serum and 1× penicillin-streptomycin in phenol red–free DMEM) and refreshed daily for 48 h. On day 2, nuclei were stained with SPY-650-DNA (1×, diluted in DMSO from a 1,000× stock) for 1 h; cells were then returned to differentiation medium and time-lapse imaging was performed on a Thunder Imager microscope (Leica Microsystems) with an HC PL APO 40×/0.95 DRY objective, capturing images every 30 min for three days.

Nuclear and cellular behavior *in vitro* during myogenic differentiation was analyzed in Fiji. Treating multinuclear aggregates as single entities, myogenic fiber formation over time was quantified as the area occupied by nuclear aggregates during differentiation.

### Myosin Heavy Chain immunofluorescence on human differentiated myoblasts

Human myoblast clones were seeded on 96-well plates (25,000 cells/well) in growth medium. At 80–90% confluence, the medium was replaced with 200 μl of differentiation medium (2% horse serum and 1% penicillin-streptomycin in DMEM) and refreshed daily for 48 h. On day 3, cells were washed with PBS, fixed with cold methanol (−20 °C, 5 min), permeabilized with 0.05% Triton X-100 in PBS (5 min, room temperature) and blocked with 100 μl/well of blocking solution (1% BSA, 2.5% goat serum, 2.5% donkey serum and 0.3% Triton X-100 in PBS). Cells were incubated with anti-MHC-I primary antibody (MF20; 1:50) for 1 h at room temperature, washed three times in PBS, and incubated with goat anti-mouse IgG, Alexa Fluor 488 (1:500) for 45 min at room temperature in the dark, followed by Hoechst nuclear staining. Imaging was performed at the ISCIII Confocal Microscopy Unit on an Operetta CLS High-Content Analysis System. MHC-I signal was quantified in the coupled software, defining plurinucleated MHC-I–positive cells as ROIs; the number of plurinucleated cells, MHC-I–positive cells and nuclei per cell, and the relative intensity, were determined.

### Statistical analysis

All statistical analyses were performed and graphs generated in GraphPad Prism 11.0. Comparisons between genotypes were assessed by one-tailed, unpaired Student’s t test, and differences were considered significant at p < 0.05.

## ABBREVIATIONS

ABE: adenine base editor
AAV: adeno-associated virus
EDMD: Emery–Dreifuss muscular dystrophy
FBS: fetal bovine serum
GFP: green fluorescent protein
L-CMD: LMNA-related congenital muscular dystrophy
MHC-I: myosin heavy chain I
NGS: next-generation sequencing
PAM: protospacer adjacent motif
sgRNA: single-guide RNA
WT: wild type

## DECLARATIONS

### Ethics approval

The human myoblast cell line used in this study (EMD2784), derived from a 3-year-old patient with *LMNA*-related congenital muscular dystrophy carrying the *LMNA* c.745C>T (p.R249W) mutation, was obtained from the MyoLine immortalisation platform, Institut de Myologie, Centre de Recherche en Myologie (Sorbonne Université–INSERM UMRS974, Paris, France), and kindly provided by Dr Gisèle Bonne under a Material Transfer Agreement. The original muscle sample was collected with written informed consent from the patient’s parents and anonymised before distribution, in accordance with French and European legislation and the EU General Data Protection Regulation. All experiments were performed in accordance with the Declaration of Helsinki.

### Consent for publication

Not applicable

### Availability of data and materials

All data generated or analysed during this study are included in this published article (and its supplementary information files).

### Authors’ contributions

M.S. acquired, analyzed, and interpreted most of the data and drafted/edited the manuscript; I.H., Da.M. and D.G.D. assisted with several experiments; Di.M. designed and performed the acquisition and analysis of the microscopy images; K.M. generated and immortalized the cell line used in this work; I.P.d.C. developed the study concept, obtained funding, coordinated and analyzed the experimental activities, and drafted/edited the manuscript. All authors have read and approved the final manuscript.

## Acknowledgements

We want to acknowledge Mario Alía, member of the Cytometry Unit from ISCIII, for carrying out the cell sorting. We thank the Genomics Unit from ISCIII for carrying out the sequencing and the loan of indices. We would also like to thank Juliana Manosalva, Clara Marín and Cristina Clemente, members of the Advanced Optical Microscopy Unit from ISCIII, for helping with processing and analysis of samples. We also thank the effort of the MyoLine immortalisation platform for immortalization of human cells from L-CMD patients. We acknowledge master thesis student Ana Quintana, Sonia Mekollaris, David Rodríguez and Salomé Ayuso for their support in the performance of different experiments.

## Declaration of generative AI in the writing process

The authors used an AI-based assistant (Claude, Anthropic) to support language editing, clarity and structuring of the manuscript. After using this tool, the authors reviewed and edited the content as needed and take full responsibility for the content of the publication.

---

**Figure S1. Transfection efficiency in human myoblasts quantified by GFP expression.** Quantification of GFP-positive cells 24 hours post-transfection across different ABE–sgRNA combinations.

**Figure S2. Adenine base editing correction of *LMNA* c.745C>T in human myoblasts.** Human myoblasts were transfected with different ABE/sgRNA combinations and DNA extracted from pools (n=2-4 replicates depending on combination) was used to amplify targeted sequenced by PCR and amplicons were subjected to deep sequencing. Graph shows the percentage of total reads obtained by the ABE-sgRNA combinations studied after CRISPResso2 analyses. Results show the percentage of the 4 main types of reads detected: wildtype, c.745T (p.R249W parental), c.743C (p.L248P bystander) and c.745T+c.743C (p.R249W+p.L248P).

