## Supplementary figures and images for "Adenine base editing correction of *LMNA* c.745C>T (p.R249W) in congenital muscular dystrophy myoblasts improves cellular phenotype while revealing deleterious p.L248P bystander effects"

### Supplemental Figures 1-2

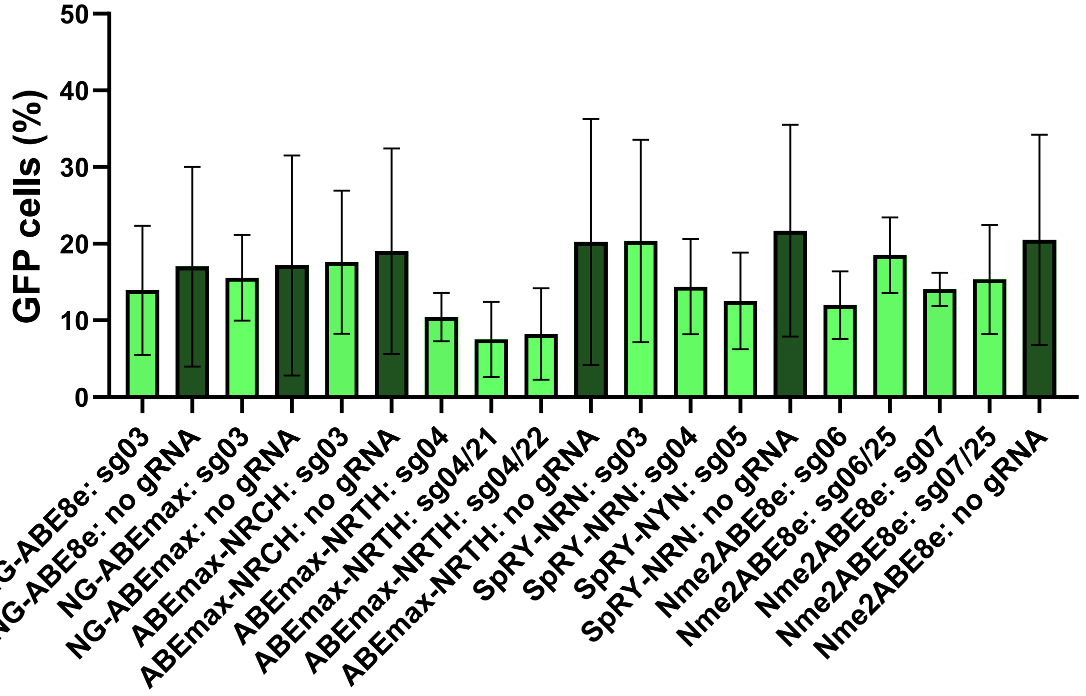

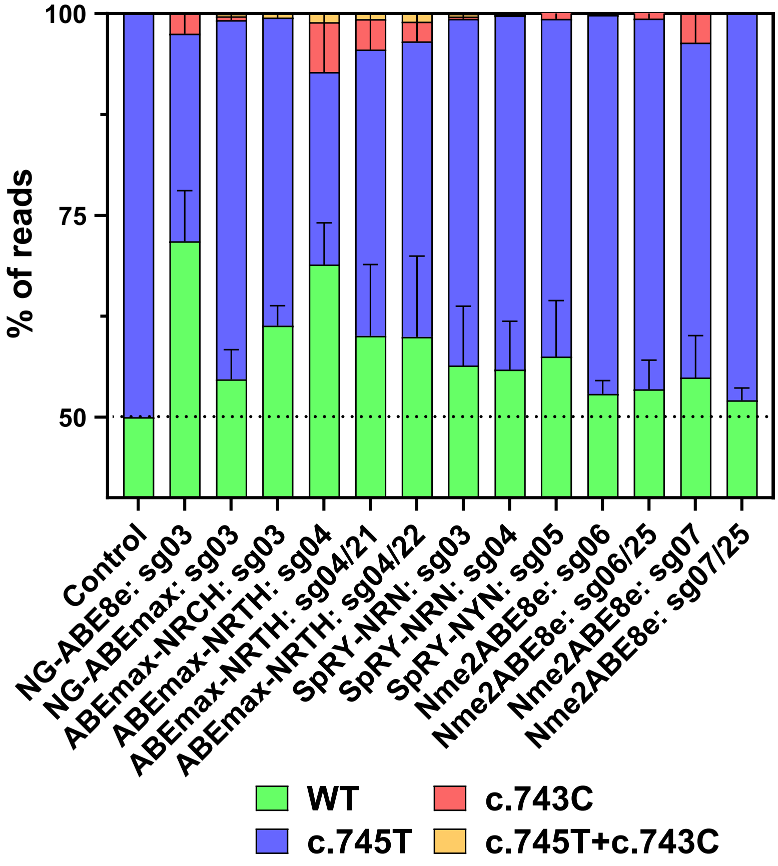
