## Supplemental Table for "Adenine base editing correction of *LMNA* c.745C>T (p.R249W) in congenital muscular dystrophy myoblasts improves cellular phenotype while revealing deleterious p.L248P bystander effects"

**Table S1. Key Resources used in this work**

| Reagent / resource | Source | Identifier |
|---|---|---|
| NG-ABE8e | Addgene | #138491 |
| NG-ABEmax | Addgene | #124163 |
| ABEmax-SpRY (pCMV-SpRY-ABE8e) | Addgene | #185671 |
| ABEmax-NRCH (pCMV-ABEmax-NRCH) | Addgene | #136923 |
| ABEmax-NRTH (pCMV-ABEmax-NRTH) | Addgene | #136922 |
| Nme2ABE8e (CMV-Nme2ABE8e) | Addgene | #189928 |
| pmaxFP-green (GFP reporter) | Amaxa / Lonza | VDF-1012 |
| Medium 199 | Invitrogen | 41150020 |
| DMEM | Invitrogen | 61965-026 |
| DMEM, no phenol red | Invitrogen | 17-205-CV |
| Fetal bovine serum (FBS) | Sigma-Aldrich | F7524-500ML |
| Horse serum | Gibco | 26050-070 |
| Penicillin/streptomycin | Lonza | DE17-602E |
| Insulin | Sigma | 91077C-1G |
| Fetuin | Life Technologies | 10344026 |
| Basic fibroblast growth factor (FGFb) | Life Technologies | PHG0026 |
| Human epidermal growth factor (EGF) | Life Technologies | PHG0311 |
| Dexamethasone | Sigma | D4902-100mg |
| Opti-MEM | Gibco | 319085-047-500mL |
| Bovine serum albumin (BSA) | Sigma-Aldrich | A7906-100G |
| Goat serum | Merck | G9023-10ml |
| Donkey serum | Merck | D9663-10ml |
| Anti-lamin A/C (E-1) | Santa Cruz Biotechnology | E-1 |
| Anti-GAPDH (14C10) | Cell Signaling Technology | #2118 |
| Anti-emerin (D3B8G) | Cell Signaling Technology | D3B8G |
| Anti-MHC-I (MF20) | DSHB | MF20 |
| Anti-mouse IgG–HRP | GE Healthcare | NA931 |
| Anti-rabbit IgG–HRP | GE Healthcare | NA934 |
| Goat anti-mouse IgG, Alexa Fluor 594 | Thermo Fisher Scientific | A-11005 |
| Goat anti-rabbit IgG, Alexa Fluor 488 | Thermo Fisher Scientific | A-11008 |
| Goat anti-mouse IgG, Alexa Fluor 488 | Thermo Fisher Scientific | A11029 |
| Hoechst | Invitrogen | 62249 |
| SPY-650-DNA | Spirochrome | SC501 |

| | | |
|---|---|---|
| µ-Slide 18-well | Ibidi | 81826 |
| 96-well plate | Corning | 353072 |
| Triton X-100 | Sigma-Aldrich | 10701922 |
| RIPA buffer | Millipore | 20-188 |
| Pierce BCA Protein Assay | Thermo Scientific | 23227 |
| Criterion TGX Stain-Free Precast Gels | Bio-Rad | 5678084 |
| Trans-Blot Turbo Transfer Pack | Bio-Rad | 1704159 |
| Pierce ECL Western Blotting Substrate | Thermo Fisher Scientific | 32106 |
| Breaking-Cas (design tool) | https://bioinfogp.cnb.csic.es/tools/breakingcas/ | |
| CRISPResso | https://crispresso.pinellolab.org/ | |
| Arivis (image analysis) | https://www.zeiss.com/microscopy/en/products/software/advanced-image-analysis.html | |
| Fiji / ImageJ | https://fiji.sc/ | |
| GraphPadPrism 11.0 | https://www.graphpad.com/ | |
